# Designing and interpreting limiting dilution assays: general principles and applications to the latent reservoir for HIV-1

**DOI:** 10.1101/018911

**Authors:** Daniel I. S. Rosenbloom, Oliver Elliott, Alison L. Hill, Timothy J. Henrich, Janet M. Siliciano, Robert F. Siliciano

**Affiliations:** Department of Biomedical Informatics, Columbia University Medical Center, New York, New York, USA; Program for Evolutionary Dynamics, Harvard University, Cambridge, Massachusetts, USA; Division of Infectious Diseases, Brigham and Women’s Hospital, Boston, Massachusetts, USA; Department of Medicine, Johns Hopkins University School of Medicine, Baltimore, Maryland, USA; Howard Hughes Medical Institute, Baltimore, Maryland, USA

**Keywords:** Dilution assay, HIV, viral outgrowth, latent reservoir, maximum-likelihood statistics, infectious unit, IUPM

## Abstract

Limiting dilution assays are commonly used to measure the extent of infection, and in the context of HIV they represent an essential tool for studying latency and potential curative strategies. To assist investigators using dilution assays, we illustrate the major statistical method for estimating the frequency of infected cells (or other infectious units) from assay results, and we offer an online tool for computing this estimate. We then recommend a procedure for optimizing assay design to achieve any desired set of sensitivity and precision goals, subject to experimental constraints. We discuss challenges involved in interpreting experiments in which no viral growth is observed and explain how using alternative measures for viral outgrowth may make measurement of HIV latency more efficient. Finally, we discuss how biological complications – such as probabilistic growth of a small infection in culture – alter interpretations of experimental results.

## 1. Introduction

HIV-1 infection persists in patients despite decades of effective antiretroviral therapy, necessitating lifelong treatment. Limiting dilution viral outgrowth assays provide an essential tool for studying this residual infection. These assays detect latently infected resting CD4^+^ T cells by probing their ability to produce virus that infects other cells in culture following cellular activation (1, 2). This latent reservoir of infected cells harbors stably integrated HIV-1 DNA coding for replication-competent virus, and as such, is believed to be the major barrier to curing HIV infection. Since replication-competent proviruses cannot be distinguished by PCR methods from the much more common defective proviruses (coding for virus that fails to replicate) (3-5), viral outgrowth remains the gold-standard marker of latent infection. Dilution assay statistics are used to estimate the frequency of latently infected cells from binary results of outgrowth experiments (6). Likelihood-based dilution assay statistics have been part of the microbiologist’s basic toolkit for nearly a century (7), and today they are indispensible to HIV-1 cure research.

Recent reports of interventions that may reduce the latent reservoir (8-14) have renewed optimism for achieving HIV-1 cure or long-term antiretroviral-free remission. Potential curative strategies are now being evaluated by how much they reduce reservoir size. It is therefore essential to improve understanding of statistics used in outgrowth assays and to establish a consistent approach for reporting the precision of the resulting measurements. Towards this goal, we present principles for the design and interpretation of dilution assays, discuss these principles in the context of HIV-1 latency, and offer a simple computational tool for estimating the frequency of latently infected cells in a sample (“infectious units per million” or “IUPM”) from assay results. The same statistical approach – and same program – can be used for a broad range of dilution assays designed to quantify viral reservoirs.

In a dilution assay, wells of a tissue culture plate are seeded with varying amounts of material from a presumed-infectious source, together with infectable reporter cells. In the viral outgrowth assay for latent HIV-1 (1,2,6), the infectious material is a known quantity of purified resting CD4^+^ T cells obtained from an HIV-1-infected donor. A very small fraction of these cells carry a latent HIV-1 genome that can give rise to replication-competent virus following cellular activation. A larger fraction of the cells carry defective viral genomes and are ignored in this analysis. Thus throughout this paper, we refer to latently infected cells that are stimulated to release replication-competent virus in the assay simply as “infected cells.”

The input cell number in each well is typically varied in geometric series of successive dilutions. Sufficient dilutions are carried out so that some wells contain no infected cells. Uninfected lymphocytes are added to each well to serve as the reporter system, so that virus released from a single cell can spread through the culture and produce a detectable level of infection. The frequency of infected cells in the original population of resting CD4^+^ T cells, as well as confidence interval around this frequency, are then estimated based on the pattern of wells in which viral outgrowth was observed. If desired, this fraction can be scaled by an appropriate number (∼10^12^ total resting CD4^+^ T cells in a human body) to estimate total latent reservoir size.

Before discussing details of assay design and statistical methods to estimate infected cell frequency, we note two issues inherent in this type of assay. First, the sensitivity – or lower limit of detection – of the assay depends on the total number of cells sampled. If *C* resting CD4^+^ T cells are distributed among all the wells, then it is impossible to distinguish between infected cell frequencies near or less than 1/*C*. Precisely at this frequency, assuming perfect experimental conditions, at least one well will show outgrowth about 63% of the time, while the remaining 37% of the time no wells will turn positive. Therefore, to reliably distinguish a frequency of 0.1 IUPM from a nonexistent infection, one would need to assay a minimum of 10 million cells, and ideally several times that number. **Section 3** further discusses lower limits of detection.

Second, the distribution of cells into wells determines the precision of the assay, its ability to distinguish between different infected cell frequencies. Precision can be described by confidence intervals: more precise assays have smaller intervals, allowing for finer-grained comparisons between experimental conditions. Figure 1 illustrates how increasing the number of wells in an assay can improve its precision. Confidence intervals are discussed more generally in **Section 2**.

**Figure 1.**
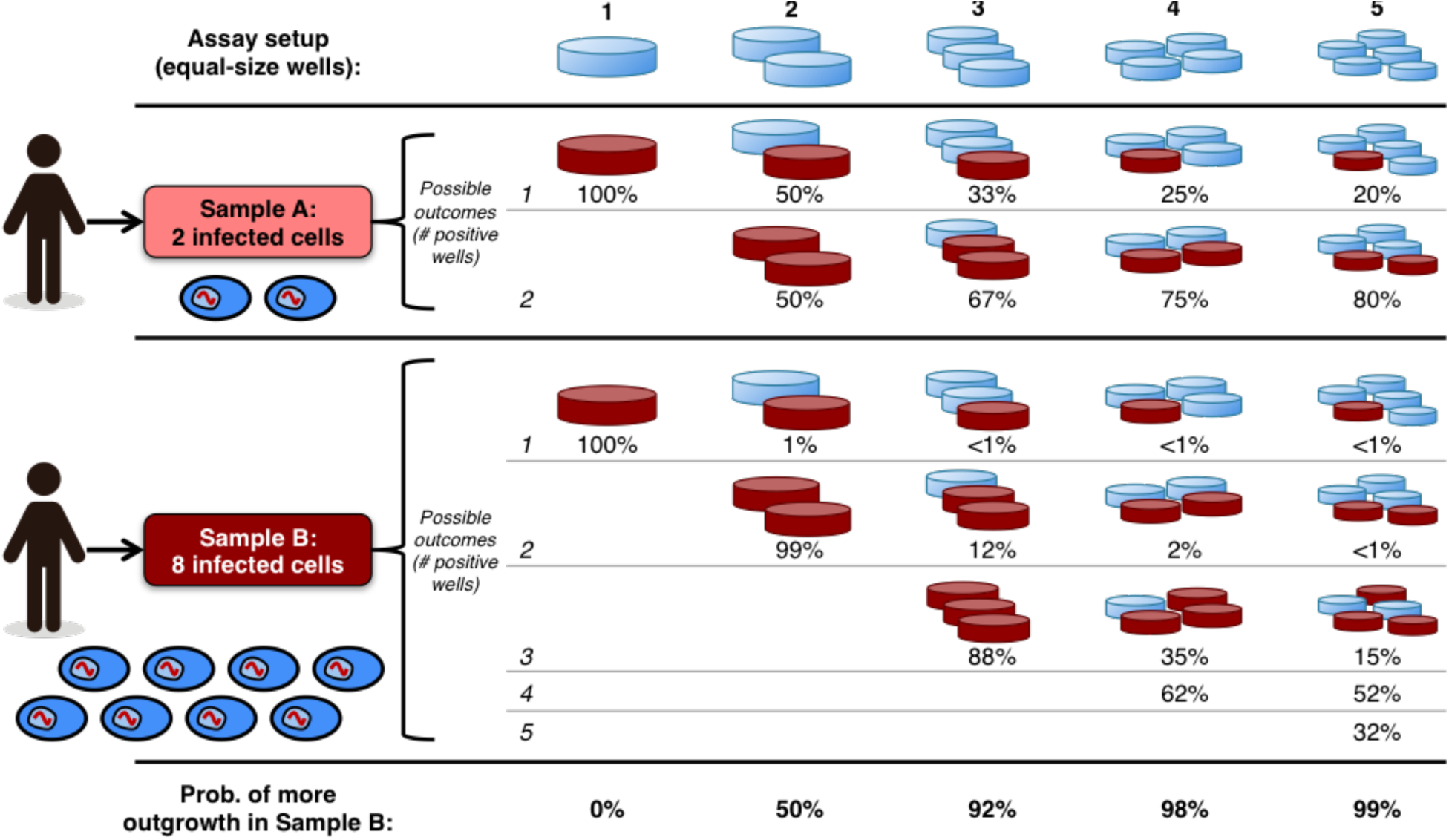
Precision of dilution assays increases with the number of wells into which a sample is divided. Consider two samples of the same size (total number of cells) collected from different donors. Sample A contains 2 latently infected cells, while Sample B contains 8 (ovals at left). Five possible assay setups are shown, distributing each sample into 1 to 5 equal wells (columns). Each assay results in a different pattern of possible outcomes for the samples, with the probability of each outcome shown in rows. The final row shows the probability that each assay yields the correct trend, that more wells turn positive in Sample B than in Sample A. Positive wells (viral outgrowth) are shown in dark red. If all cells from a sample are deposited into a single well (first column), then it is impossible to distinguish between the two samples using a binary readout: the extra infected cells in Sample B are “redundant,” and each sample will show outgrowth. If the samples are divided among more wells (second through fifth columns), then the infected cells are less likely to appear redundantly in the same well, meaning that the two samples are more likely to show different outgrowth patterns. In this particular case, each sample must be divided into at least four equal wells in order to obtain the correct trend at least 95% of the time.

Sensitivity requires many cells, and precision requires distribution into multiple wells. Meeting both goals presents experimental challenges. Next we present likelihood statistics to estimate infected cell frequency, and we recommend an approach for optimizing experimental design to achieve any desired set of sensitivity and precision goals. While our discussion focuses on HIV-1 infection, the same approach applies to any setting where frequency of infected cells is measured.

## 2. Likelihood statistics provide a flexible approach for estimating infected cell frequencies

The most common method for analyzing limiting dilution assays involves maximum likelihood estimation of infected cell frequencies. This general method can be used for any experiment in which the observed outcome depends on one or more unknown parameters in a probabilistic way. Here, the only unknown parameter is the infected cell frequency, and the maximum likelihood method provides both a central estimate and confidence interval for this frequency. If the infected cell frequency lies within the assay’s *limits of detection*, then the maximum likelihood estimate is expected to be quantitatively meaningful, falling between but not equal to the extreme values 0 IUPM and 1,000,000 IUPM. This estimate furthermore will be precise – with confidence intervals (CI) smaller than a specified size – for frequencies within the assay’s (smaller or coinciding) *limits of quantification*. These concepts are further explained in Figure 2. In general, within the limits of quantification, the size of the CI depends on the experimental setup, and so CIs must be reported explicitly to allow comparison of infected cell frequency over time or among patients.

**Figure 2.**
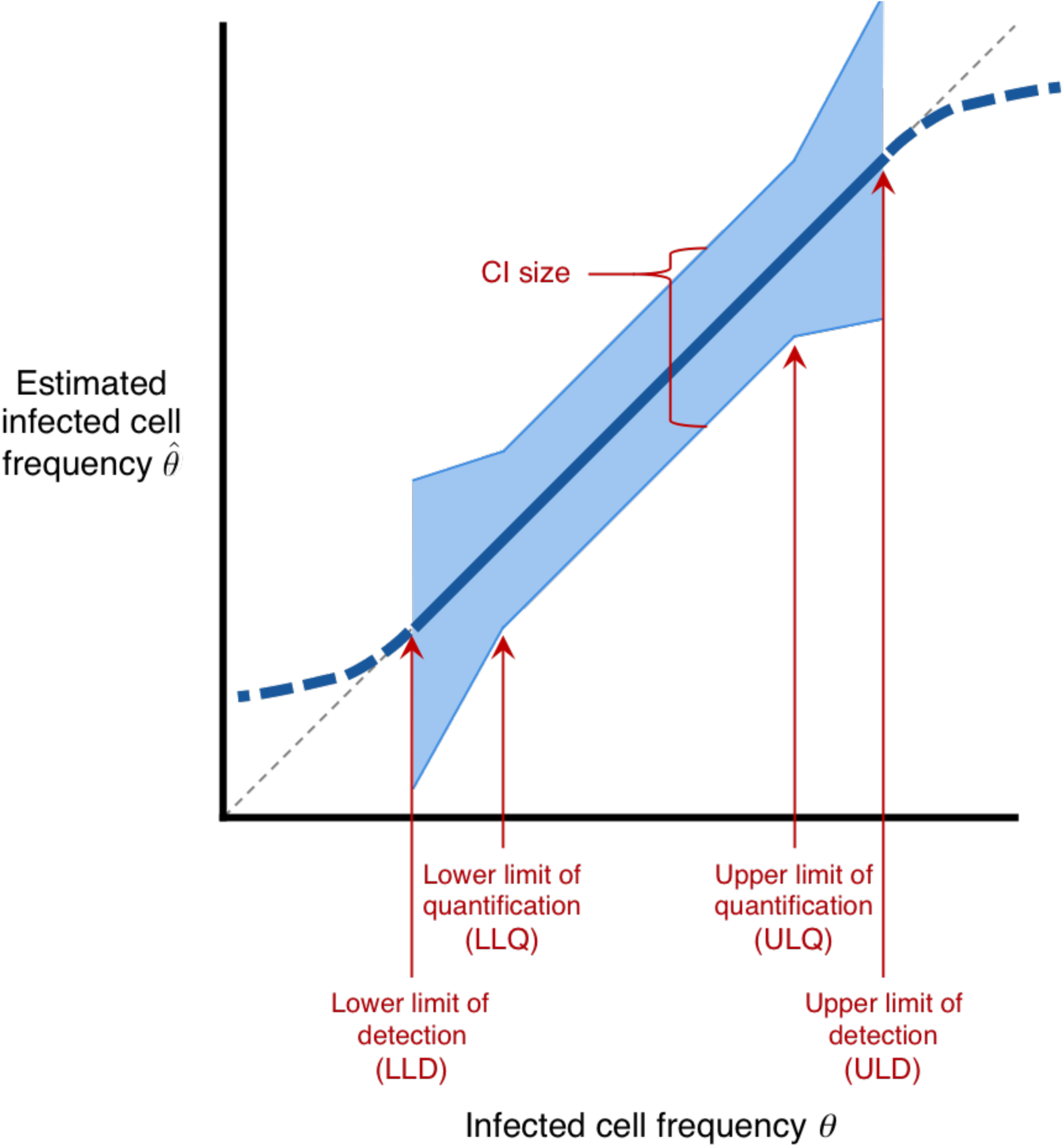
Schematic for describing quality of maximum-likelihood estimates of infected cell frequency. The thick blue line plots the typical value (median or geometric mean) of the measured infected cell frequency 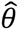 (y-axis) versus true frequency *θ* (x-axis), for an idealized experimental setup; this line is dashed outside the limits of detection for the assay. These limits are defined for a significance level α (e.g., 0.05). Below the **lower limit of detection (“LLD”)**, at least α of experiments result in all wells being negative. Above the **upper limit of detection (“ULD”)**, at least α of experiments result in all wells being positive. In an accurate assay, the estimate tracks the diagonal line 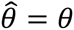 (light gray dashed line) between the limits of detection. The blue shaded region plots the **confidence interval (CI)**, within which the middle (1 – *α*) of estimates fall. Between the **lower and upper limits of quantification (“LLQ", “ULQ”)**, the CI is smaller than a specified size, *and* a fraction less than *α* of experiments result in all-positive or all-negative wells. Note that it is possible for LLD = LLQ and/or ULD = ULQ.

The basic principles of maximum likelihood analysis for dilution assays have been well understood since the work of Fisher (7) (see also (15)). These principles have also been extended to a Bayesian framework, which explicitly accounts for how prior knowledge alters interpretation of experimental results (16). A study by Meyers et al. (6) presented a complete analysis of maximum likelihood estimates and confidence intervals resulting from all possible experimental outcomes of a particular fivefold limiting dilution assay design with two replicates per dilution. This limiting dilution scheme was used in early studies of HIV-1 latency (1) for a particular reason. The low frequency of latently infected cells necessitates sampling a large number of cells, but blood volume concerns limit the number of replicate wells that can be done with higher input cell numbers. This assay design is widely used in standard assays of the latent reservoir (17), but may not be optimal for all experimental situations. Specifically, infected cell frequencies may lie outside this assay’s limits of detection or quantification, and CIs may be too large to address particular research questions. Fortunately, custom assays can be designed to satisfy any desired limits and CI sizes; in Box 1, we present a simple procedure for doing so. Table 1 shows several recommended designs resulting from this procedure.

**Table 1.** Recommended dilution assay designs satisfying a range of experimental requirements as provided in Box 1, using a significance level *α* = 0.05, corresponding to 95% confidence intervals. The dilution series commonly used in HIV-1 latency studies (1,6,17) is shown on the first row. Simulated performance of the two shaded rows is shown in Figure 3. Since decreasing the LLD and CI size each involves using more cells according to the procedure in Box 1, these two requirements are decreased together in this table.

| Experimental requirements |  |  |  | Resulting assay design |  |  |  |  |
| --- | --- | --- | --- | --- | --- | --- | --- | --- |
| Limits of detection, quantification (IUPM) | | | CI size ($\log_{10}$) | # different cell input numbers (obtained by five-fold dilution) | Largest well (input cells) | Smallest well (input cells) | # replicates (largest to smallest) | Total cells (millions) |
| LLD | LLQ | ULQ = ULD |  |  |  |  |  |  |
| 1.20 | 1.60 | 800 | < 1.5 | 6 | $1 \times 10^6$ | 320 | 2 each | 2.5 |
| 0.48 | 1.60 | 800 | < 1.0 | 6 | $1 \times 10^6$ | 320 | 5 each | 6.2 |
| 0.13 | 1.60 | 800 | < 0.5 | 5 | $1 \times 10^6$ | 1600 | 18 each | 22 |
| 0.20 | 0.50 | 12.5 | < 1.5 | 4 | $3.2 \times 10^6$ | $2.56 \times 10^4$ | (4, 2, 2, 3) | 14 |
| 0.15 | 0.50 | 12.5 | < 1.0 | 4 | $3.2 \times 10^6$ | $2.56 \times 10^4$ | 5 each | 20 |
| 0.05 | 0.50 | 12.5 | < 0.5 | 3 | $3.2 \times 10^6$ | $1.28 \times 10^5$ | 18 each | 71 |
| 0.04 | 0.10 | 12.5 | < 1.5 | 5 | $1.6 \times 10^7$ | $2.56 \times 10^4$ | (4, 2, 2, 2, 3) | 72 |
| 0.03 | 0.10 | 12.5 | < 1.0 | 5 | $1.6 \times 10^7$ | $2.56 \times 10^4$ | 5 each | 100 |
| 0.01 | 0.10 | 12.5 | < 0.5 | 4 | $1.6 \times 10^7$ | $1.28 \times 10^5$ | 18 each | 360 |
| 0.0080 | 0.02 | 12.5 | < 1.5 | 6 | $8 \times 10^7$ | $2.56 \times 10^4$ | (4, 2, 2, 2, 2, 3) | 360 |
| 0.0006 | 0.02 | 12.5 | < 1.0 | 6 | $8 \times 10^7$ | $2.56 \times 10^4$ | 5 each | 500 |
| 0.0002 | 0.02 | 12.5 | < 0.5 | 5 | $8 \times 10^7$ | $1.28 \times 10^5$ | 18 each | 1,800 |

Formally, maximum likelihood estimation of infected cell frequency relies on four assumptions justifying the use of Poisson distributions to approximate the number of infected cells in each well: First, that cells in the sample are drawn from a much larger total population (e.g., 50 million cells sampled from ∼10^12^ total resting CD4^+^ T cells in the body). Second, that the infected cell frequency is small (e.g., frequency less than 1 in 100 ensures approximation error less than 0.5%). Third, that infected cells are distributed randomly among all replicate wells. Finally, that any well containing one or more infected cells is guaranteed to test positive.

Together, these assumptions constitute the “single-hit Poisson model.” In this model, the probability that a well with *c* cells tests positive, given an infected cell frequency *θ*, is 1 – *e*^–*cθ*^. The number of positive wells of a given input cell number *c* follows the binomial distribution specified in Figure 3. When wells of different input cell numbers *c*_1_, *c*_2_, *c*_3_, … are used, the results from each are independent, and so the probability of an observed experimental result is simply the product of individual binomial probabilities. The program IUPMStats (http://silicianolab.johnshopkins.edu) uses this calculation to estimate the infected cell frequency *θ*, as well as 95% confidence intervals, from user-specified assay results. Figure 3 shows the performance of this method for two of the assay designs proposed in Table 1.

#### Box 1: Procedure for designing a limiting dilution assay

In designing an assay, the experimenter must choose a range of input cell numbers and the number of replicate wells to set up at each input cell number. This choice determines the lower and upper limits of detection (*LLD*, *ULD*), lower and upper limits of quantification (*LLQ*, *ULQ*), and confidence interval (CI) size *ε* (expressed as a number of base-10 logs). The limits of detection determine the range in which the assay rarely produces all-positive or all-negative results. Between the limits of quantification, the assay furthermore can distinguish infected cell frequencies up to a range of *ε* logs. Figure 2 further explains these concepts. Below, *z*_*α*_ refers to the critical normal distribution value corresponding to significance level *α* (e.g., *Z*_0.05_ = 1.96, *Z*_0.01_ = *2*.58).

The following procedure is guaranteed to produce an assay design with specified limits *LLD* ≤ *LLQ* < *ULQ* ≤ *ULD* and maximum CI size *ε*, for a significance level *α*, aiming to use the fewest wells necessary (derivation in **Supplementary Information**).

1. Select desired limits and maximum CI size. Experimental requirements may constrain feasible values:
  A. If *C* cells are available to distribute across all wells, then the lower limit of detection is constrained: *LLD* ≥ ln (1/*α*) /*C*.
  B. If the experimenter wishes to perform at most *n* replicate wells per input cell number, then the CI size is constrained: 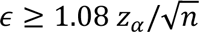.
  C. If the experimenter wishes to use only *d* distinct input cell numbers, that constrains the width of the region between the limits of quantification: *ULQ/LLQ* ≤ 5^*d*^−1.
  D. If *C* cells are available *and* the experimenter wishes to perform at most *n* replicate wells per input cell number, then in addition to the above constraints on *LLD* and *∈*, we have *LLQ* ≥ 1.99 *n/C*.
  E. If *C* cells are available *and* the experimenter wishes to guarantee a CI size of at most *ε*, then in addition to the above constraint on *LLD*, we have 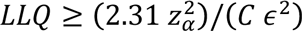.
2. To guarantee that the CI is at most *ε* logs for infected cell frequencies within the limits of quantification, choose a limiting dilution series where:
  A. Maximum input cell number *c*_max_ is at least 1.59/*LLQ* cells,
  B. Minimum input cell number *c*_min_ is at most 1.59/*ULQ* cells,
  C. Each input cell number must differ by no more than a factor of 5 from the previous one,
  D. Each input cell number has at least 1.16 × *z*_*α*_/*∈*)^2^ replicates.
3. To guarantee the desired limits of detection, further require:
  A. The total number of cells across all wells is at least ln (1/*α*)*LLD*,
  B. The smallest input cell number is at most ln (1/(1 − α^1/*n*^))/*ULD*, where *n* is the number of replicate wells of that cell number.

Often, the dilution series provided by Step 2 will also satisfy the conditions in Step 3 and may provide limits of detection even farther apart than required. In this case, nothing needs to be changed. If, however, the dilution series does not meet condition 3A, wells of any input cell number can be added to make up the difference in total cells. To meet condition 3B, either the number of replicates for the smallest well may be increased, or wells of a smaller input cell number may be added. It is possible for this condition to require a few additional wells beyond that specified in the experimental constraints in Step 1.

This procedure may yield input cell numbers that are impractically large for loading into wells. From the standpoint of ensuring the required limits and confidence interval sizes, it is always acceptable to replace a well containing many input cells by multiple wells containing fewer input cells each, keeping the same total number of cells. For instance, replacing two wells of 10 million cells apiece by ten wells of 2 million cells apiece can only serve to increase precision of the maximum likelihood estimate. If it is experimentally convenient to have all wells with the same input cell number, then this principle can likewise be used to do so.

**Figure 3.**
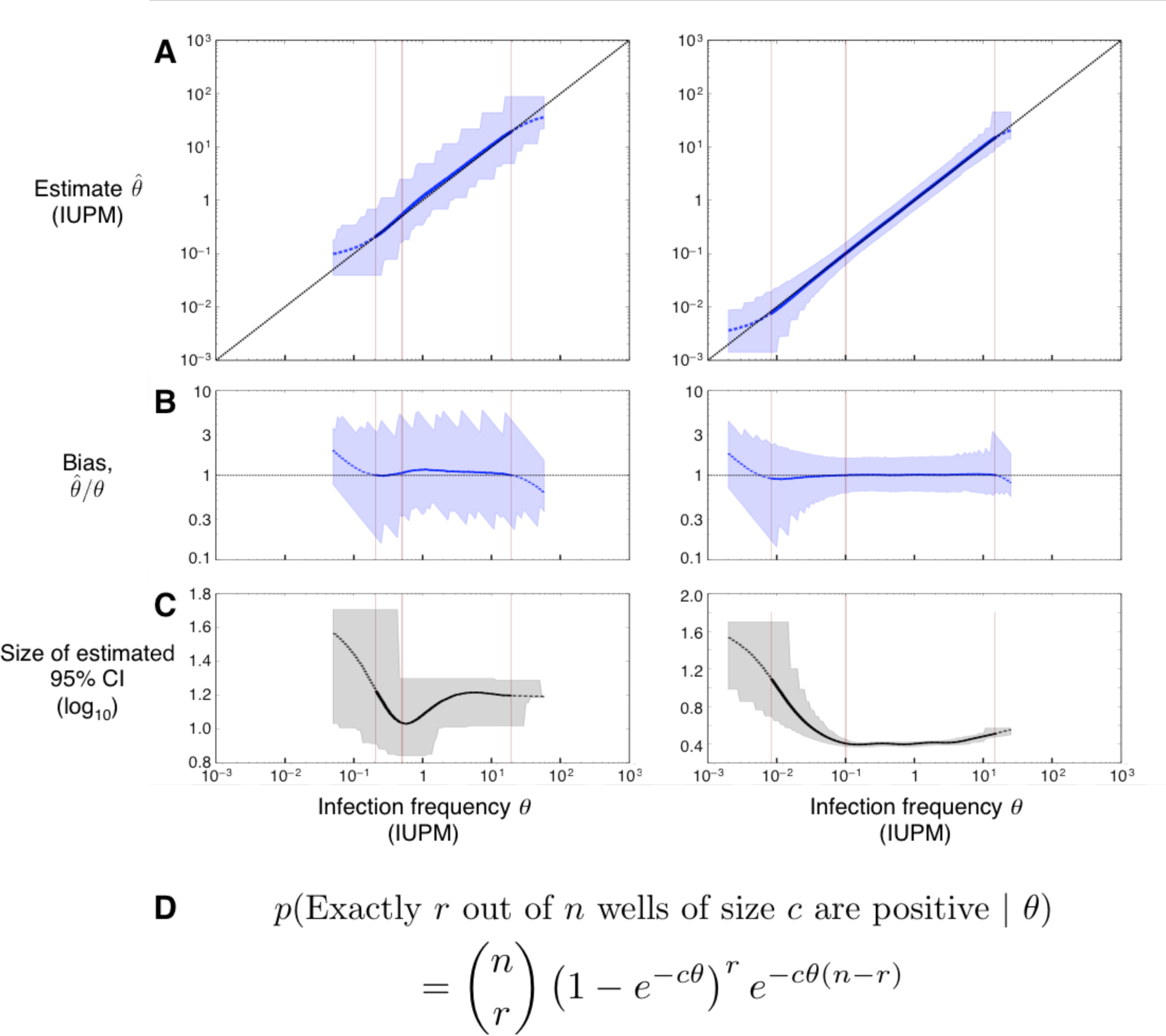
Performance of the maximum likelihood estimator in simulations, using the two assay designs highlighted in Table 1. Left column: assay on row 4 (CI < 1.5 logs); right column: assay on row 9 (CI < 0.5 logs). **A)** Maximum likelihood estimate 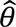 and 95% confidence interval plotted in blue; the diagonal line shows the case of a perfect unbiased estimator. **B)** Bias 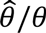 plotted in blue; the horizontal line at 1 shows the case of a perfect unbiased estimator. **C)** Size of the estimated 95% confidence interval plotted in black (note different y-axis scales). **D)** Binomial probability expression used to estimate infection frequency, assuming *n* replicate wells of *c* cells apiece. According to this expression, the probability that all wells are negative equals *e*^*−cθ*^, where *C* is the total number of cells across all replicate wells. **Panels A – C** plot the actual infection frequency *θ* used in simulations on the x-axis. Each point on the curves is the geometric mean (A, B) or arithmetic mean (C) of 20,000 replicate simulations using the same *θ* (step size 0.025 logs). Curves are solid where < 5% of simulated assays yield all-negative or all-positive results, dashed at 5% – 50%, and not shown at > 50%. Blue (A, B) or gray (C) shaded regions show the middle 95% of simulations; jaggedness results from the discrete nature of the dilution assay. Left to right in each panel, the thin vertical lines show the LLD, the LLQ, and the ULQ = ULD. Note that the assay shown at right is more sensitive (20-fold lower LLD), more precise (narrower shaded regions in Panels A and B, smaller CI in Panel C), and more accurate (curve in Panel A better tracks the diagonal; curve in Panel B better tracks the horizontal).

## 3. Dealing with all-negative data: absence of evidence is not evidence of absence

If all wells in an assay test negative for viral growth, then the maximum likelihood estimate of infected cell frequency is zero. Yet this is not a useful estimate; it is possible for some of a patient’s cells to be infected (*θ* > 0), but for none of these infected cells to be sampled. The probability of this all-negative outcome follows from the equation in Figure 3D.

To avoid this outcome, the procedure in Box 1 prescribes an assay design based on a specified lower limit of detection (LLD). The total number of cells *C* is chosen so that the probability of finding an all-negative result is small (e.g., *α* = 0.05) if *θ* = LLD. As *θ* falls below the LLD, the probability of an all-negative result increases, making it difficult to distinguish between infection frequencies below this limit. For significance level *α*, the LLD equals ln (1/*α*)/*C*. For example, if we want to be nearly certain (> 95%) of detecting infection at frequencies as low as 0.01 IUPM, then 300 million cells are needed.

When an all-negative result does occur, however, some estimate for infected cell frequency is needed. Under conservative assumptions (uniform Bayesian prior on infected cell frequency between 0 and 1,000,000 IUPM), and for sufficiently many total cells *C* (> 500), this result implies, with probability (1 − *α*), that the infected cell frequency falls below the LLD The LLD, then, is a conservative upper bound for the estimated frequency. If a more central estimate is needed, then under the same assumptions, there is a 50% chance that infected cell frequency is less than ln (2)/*C*; this value is called the “posterior median” estimate. For infected cell frequencies near this value, longitudinal samples may switch between testing all-negative and having some wells positive, even if the true frequency remains constant over time.

It is important not to overinterpret these statements: The LLD is not a “hard cutoff” below which the assay is useless. Furthermore, it is possible for the maximum likelihood estimate 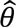 to be below the LLD even if some wells are positive. One should therefore take care when drawing conclusions from negative results, or when comparing negative results to other experiments where 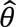 is near the LLD. In general, formal hypothesis testing is required when comparing estimates near limits of detection.

It is likewise important to stress that the absence of evidence is not evidence of absence: The most sobering recent demonstration of this principle is the case of the “Mississippi Child,” who underwent combination antiretroviral therapy for the first 18 months of life after being infected with HIV-1 *in utero*. Despite subsequent interruption of therapy, viral rebound did not occur immediately and repeated outgrowth assays found no replication-competent virus, raising tentative hopes that early aggressive treatment succeeded in preventing establishment of the latent reservoir (11). These hopes were dashed when the virus rebounded after 28 months off therapy (14). Four separate viral outgrowth assays were done throughout the interruption, with a total of 66 million resting CD4^+^ T cells analyzed. Since the latent reservoir decays very slowly (2, 18), it is reasonable to combine negative assay results and estimate that, with 95% probability, there were fewer than ln 1 0.05 66 = 0.045 infectious units per million cells (14). Consistent with the observed viral rebound, mathematical modeling of infection dynamics suggests that even this low frequency is insufficient to make cure likely (19).

## 4. New dilution assays to measure viral reservoir – better sensitivity for equal cell input?

Given the time, labor, and large cell input required for sensitive measurement of latent HIV infection, there is great interest in the development of assays to quantify HIV reservoirs more efficiently. One recent innovation is the Tat/Rev Induced Limiting Dilution Assay (TILDA), which measures cell-associated multiply-spliced HIV-1 RNA (msRNA) transcription in serial dilutions of maximally stimulated CD4^+^ T cells (20). As with outgrowth assays, TILDA uses dilutions of a known number of cells distributed across replicate wells and gives a binary readout. Consequently, the same maximum likelihood approach – and the IUPMStats program – can be used. Unlike outgrowth assays, however, the use of msRNA measurement in TILDA eliminates the need for infectable reporter cells and permits a rapid turnaround time (days, versus weeks for outgrowth assays). Because TILDA measures an upstream process that is necessary (21-23), but not sufficient for assembly of replication-competent virus, it is not surprising that the number of msRNA-producing units per million (“msRUPM”) exceeds outgrowth-based IUPM in patients receiving antiretroviral therapy (20). As a result, TILDA (and other assays of upstream processes) will appear more sensitive than outgrowth, in the sense that fewer total input cells are needed to detect at least one positive well. Yet it remains to be shown whether “msRUPM” or related measurements are clinically useful proxies for the size of the replication-competent latent reservoir, given that this assay may detect mRNA production from cells with defective proviruses (3).

## 5. When the assumptions fail

As described in Section 2, the statistical framework for estimating infected cell frequency relies on several assumptions. One way in which these assumptions may fail is for each infected cell to cause a positive result with probability less than one. There are several ways that a latently infected cell, capable of releasing replication-competent virus upon stimulation, may nonetheless fail to trigger outgrowth. For instance, variation in host gene expression may stall initiation of HIV transcription (3, 24). Furthermore, even if transcription starts, noisy transactivation feedback circuits may fail to cause elongation of viral transcripts (3, 25-27). Some stimuli increase HIV-1 gene expression but do not result in packaging and export of virus particles (13, 28). Finally, the virions produced may fail to establish an exponentially growing infection (19). These types of failure, however, do not change the basic statistical framework of the assay, only its interpretation. In each scenario described, functionally intact provirus escapes detection in the outgrowth assay. In this light, the quantity being estimated in the assay is a weighted sum of outgrowth probabilities, considering all cells carrying potentially inducible replication-competent proviruses. For example, a measured frequency 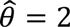 IUPM is consistent with a population where one per million cells have 100% chance of outgrowth, while an additional ten per million have 10% chance of outgrowth. The “infectious unit” of IUPM therefore refers to a statistical quantity causing viral outgrowth under laboratory conditions which may consist of many infected cells. This interpretation of IUPM follows common usage in the study of infectious diseases, which treats “infectious unit” as the quantity of pathogen causing a single infection *in expectation* (29), in contrast to the less frequent usage treating it as the *minimum* quantity capable of causing infection (30). Sensitivity of IUPM measurements to assay design further attests to the chance nature of outgrowth. While little discussed, this sensitivity is tacitly documented in the literature by experimental protocols that recommend longer periods of monitoring for outgrowth if surprisingly few wells turn positive (17).

A second type of failure arises if infected cells are not distributed randomly among wells, which may occur, for instance, if tissue from an infected donor is not fully dissociated. In this case, infected cells may cluster in the same well, causing underestimates in infected cell frequency. Poor mixing may also lead to counterintuitive assay results – e.g., positive results at small input cell numbers but not at large numbers. Tests for goodness of fit to the single-hit Poisson model can flag such outcomes (6); an approximate *χ*^2^ test is included for this purpose in IUPMStats. Standard experimental procedures are designed to ensure random mixing of cells and avoid these outcomes.

Third and most troubling is the possibility that outgrowth depends on the number of infected cells in a non-independent way. There are several mechanisms by which multiple infected cells occupying the same well may interact synergistically to increase the probability of detectable outgrowth. HIV, like both simian and murine retroviruses, can saturate host restriction factors at high viral concentrations, increasing the probability that a cell is successfully infected (31). Additionally, sustained viral production depends on intracellular concentration of the viral protein Tat (25, 26), which may be able to cross cell membranes and reach critical concentrations more readily when multiple infected cells are present (32). If these concentration-dependent mechanisms occur under experimental conditions, then estimated infected frequency 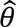 may increase if input cells are plated at higher concentrations. In this case, a more complex “multi-hit” model would be needed to estimate the size of the latent reservoir.

Accurate measurement of the latent infection in HIV continues to be a major challenge. As the field continues to develop therapies to reduce latency, principled design and interpretation of dilution assays will become increasingly important in prioritizing these therapies for clinical trial. We hope that our online tool (http://silicianolab.johnshopkins.edu) can assist the field in this task.

## Acknowledgements

We thank Joel Blankson, Christine Durand, Evelyn Eisele, Ya-chi Ho, Jun Lai, Greg Laird, Adam Longwich, Afam Okoye, Deborah Persaud, María Salgado Bernal, Carrie Ziemniak, and Michelle Kim Zinchenko for helpful discussions about assay design and for testing the IUPMStats program. This work was funded by NIH grants R01GM109018 (DISR), R01CA185486 (DISR), DP5OD019851 (ALH), and 1K23AI098480-01A1 (TJH) and an amfAR Research Consortium on HIV Eradication grant (TJH).

## Supplementary Information

### Justification of assay design procedure

The design procedure in Box 1 of the Main Text guarantees specified limits of detection and quantification, for a desired significance level *α* and confidence interval width *ε*. As in the Main Text, *θ* is the actual infected cell frequency and 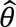 is the maximum likelihood estimate. The assay design is given by the vectors 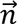 and 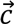, where there are *n*_*i*_ replicate wells of *c*_*i*_ input cells apiece, listed in order of decreasing *c*_*i*_. The total number of cells in the assay is 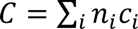 and *c*_max_ and *c*_min_ are the largest and smallest cell input numbers, respectively.

The **lower limit of detection** (*LLD*) is defined as the infected cell frequency such that all wells of the assay are negative with probability α. This result occurs if none of the sampled cells are infected, which occurs with probability (1 − *θ*)^*c*^, which is approximately *e*^*−cθ*^ for small *θ*. The required number of cells to ensure a desired lower limit of detection is therefore *C* = ln (1 − *α*)/*LLD*. A larger *C* is acceptable, as it would serve to decrease the lower limit further.

The **upper limit of detection** (*ULD*) is defined as the infected cell frequency such that all wells of the assay are positive with probability α. This result occurs if at least one sampled cell from each well is infected, which occurs with probability 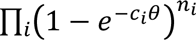. A more concise sufficient condition for ensuring that an upper limit exceeds a desired value can be given by providing that all wells of the smallest cell input number are positive; in other words, by solving for *c*_min_ in the equation α = (1 − *e*^−*c*_min_*θ*^)^n^, where *n* is the number of replicate wells with *c*_min_ cells. The result is *c*_min_ = ln(1/(1 − *α*^1/*n*^))/*ULD*. A smaller *c*_min_ is acceptable, as it would serve to increase the upper limit further.

For infected cell frequencies between the **lower and upper limits of quantification** (*LLQ*, *ULQ*), the confidence interval at significance level *α* has a width less than *ε*. The expected Fisher information on ln (*θ*) can be used to estimate confidence interval width ^1,2^. The information provided by *n* wells with *c* cells each is *g*(n, c; *θ*) = *n*(*c*^2^ *θ*^2^)/*e*^*cθ*^ − 1), and the total information for the experiment is the sum of information across all wells, 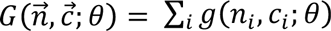. The variance in the estimate of ln (*θ*) is the inverse of this total information *G*. It follows that the expected confidence interval size, expressed as a number of base-10 logs, is equal to

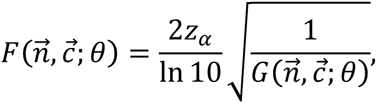

where *z*_*α*_ is the critical normal distribution value for significance level *α* (e.g., *z*_0.05_ = 1.96). The goal for assay design is to ensure that the CI size 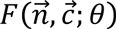 remains below *ε*, for all frequencies in the range *LLQ < θ < ULQ*. Since there may be many well sizes, it is complex to optimize based on this criterion in general. A simpler sufficient condition may be obtained by considering the confidence intervals that would be provided by a single cell input number individually:

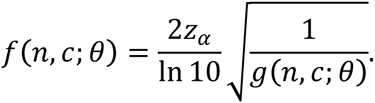

Here, the lowercase *f* denotes CI size provided by wells of just one cell input number, as opposed to capital *F* for the entire assay with multiple cell input numbers. The cell input number that minimizes this CI size, holding other parameters constant, is *c* ≈ 1.59/*θ*. To ensure *f*(*n, c; θ*) ≤ *ε*, holding *c* and *θ* constant, the number of replicates must be *n* ≥ 1.16 × (*z*_*α*_/*∈*)^2^. To ensure that the CI size is below *ε* at two frequencies *θ*_1_ and *θ*_2_, then clearly the procedure can be repeated to yield *n*_1_ wells of size *c*_1_ (ensuring *f*(*n*_1_, *c*_1_; *θ*_1_ ≤ ∈) wells of size *c*_2_ (ensuring *f*(*n*_2_, *c*_2_; *θ*_2_ ≤ ε). If *θ*_1_ and *θ*_2_ differ at most six-fold, then it is also the case that *F*([*n*_1_, *n*_2_], [*c*_1_, *c*_2_]; *θ*) < ε for all *θ* between *θ*_1_ and *θ*_2_; this fact can be verified by noting that the derivative of *F* with respect to *θ* is nonzero within this interval, provided that *c*_1_ and *c*_2_ differ at most six-fold. A CI smaller than *ε* can be ensured for an arbitrarily wide interval by using multiple dilutions such that *c*_*i*_ and *c*_*i*+1_ differ at most six-fold for all *i*. In Box 1 in the Main Text, we make a slightly more conservative recommendation (five-fold instead of six-fold dilution) to account for the fact that the actual confidence interval estimate can exceed the expected value.

The experimental constraints given in Step 1 of **Box 1** follow directly from the relationship between *C* and *LLD*, the relationship between *n* and *∈*, the cell input number minimizing CI size at frequency *θ*, and the requirement of at most five-fold dilution.

